## supplemental material for "Virus-encoded Shemin pathway highlights the importance of tetrapyrrole metabolism during host infection"

**Figure S1**

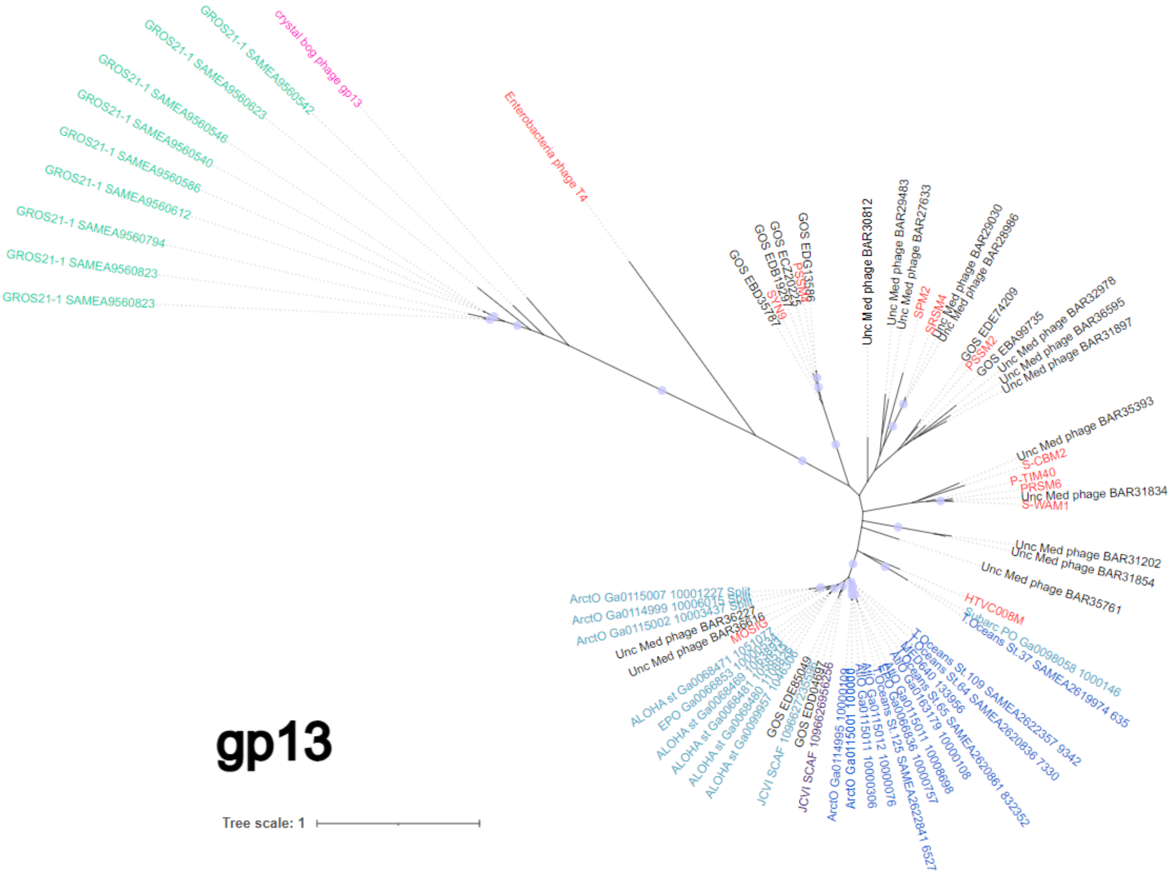


**Figure S1: Phylogenetic analyses based on gp13 protein sequences**. Maximum likelihood phylogenetic trees for gp13. Annotation in the phylogenetic tree represent gp13 source contigs/phages. Cultured phages are coloured red; black names denote metagenomic contigs of uncertain origin. The CB_2 phage chosen for experimental characterization is coloured pink. The previously characterized cassette containing a *hemO_pcyX* (Ledermann et al., 2016) is marked in purple. Metagenomically retrieved contigs form this project are colour coded according to the bars in Figure 2. Circles represent bootstrap values >0.9. The scale bar indicates the average number of amino-acid substitutions per site.

**Figure S2.**

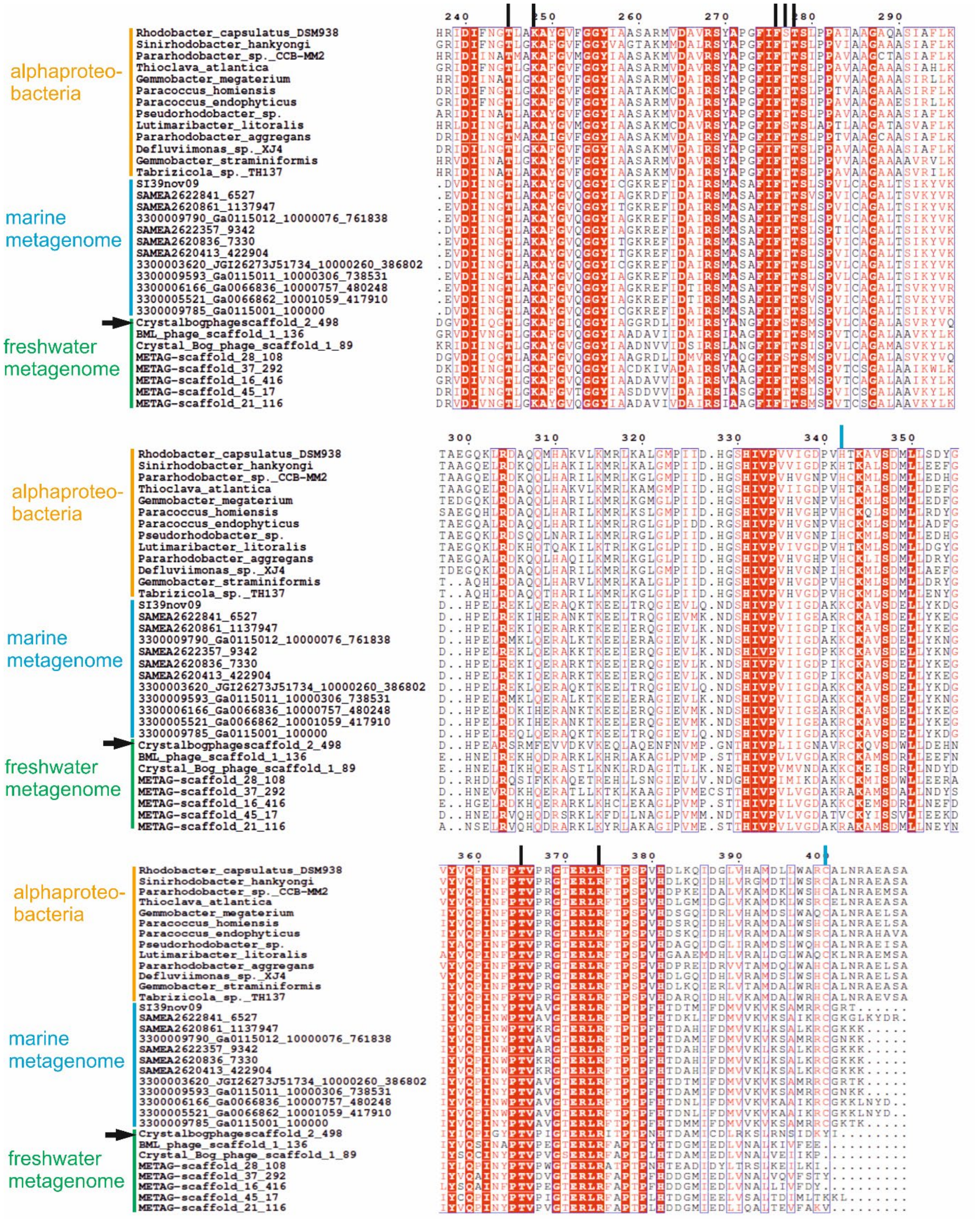


**Figure S2. Partial amino acid sequence alignment of AlaS sequences from alphaproteobacteria and viral metagenomes revealed conserved catalytic residues.** Reference sequences (shown in yellow) from the NCBI database were aligned with sequences from multiple sampling sites (blue and green) using Clustal Omega and ESPript 3. Highly conserved amino acids are highlighted in a red square with white lettering, similar amino acids are represented by a white box with red lettering, and non-similar amino acids are indicated by black lettering. Catalytic important amino acids of RcA, previously characterized in (Astner et al., 2005), are denoted by black lines at the top. Heme axial ligands of *Caulobacter crescentus* AlaS are indicated with blue lines at the top (Ikushiro et al., 2018). Marked with an arrow is the sequence used in this publication for characterization.

**Figure S3.**

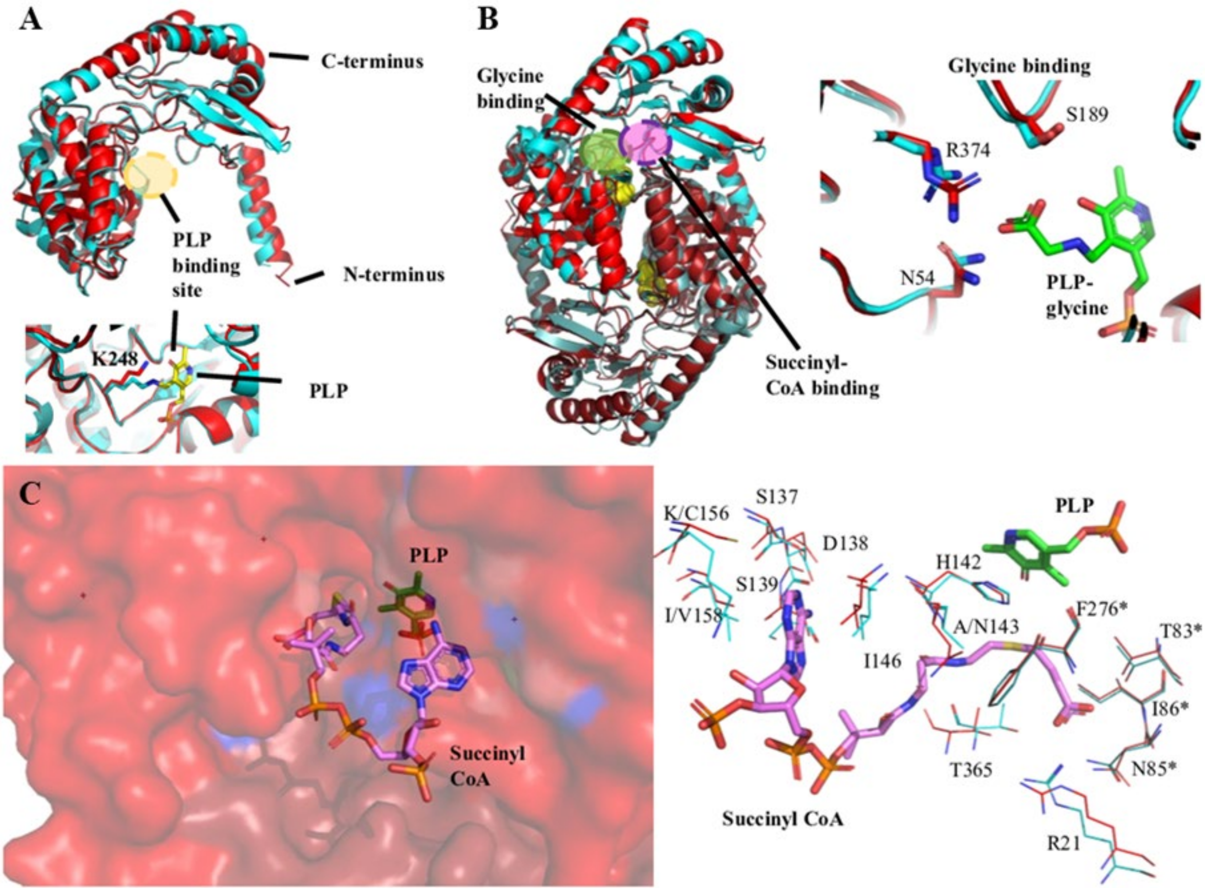


**Fig. S3. Structural modelling of vAlaS revealed high similarity to bacterial AlaS.**
**A.** Overlay model of monomeric AlaS from *Rhodobacter capsulatus* (RcA, blue, RCSB PDB: 2bwn) and the model of vAlaS predicted via AlphaFold (red), shows high structural similarity. The cofactor binding site of PLP is indicated with a yellow circle. A magnified view shows binding of PLP (yellow) onto amino acid K248 (sticks in blue/red) forming a Schiff's base linkage. **B.** Homodimeric view of vAlaS with RcA overlay, indicated are glycine binding (green circle) and succinyl-CoA binding (purple circle) and an enhanced view of the three important amino acid residues of the glycine binding (N54, S189, R374) onto the PLP-glycine intermediate (green chemical structure). **C.** The substrate channel for succinyl-CoA binding is visualized, with the ribose moiety outside the enzyme (purple) and PLP (green) inside (left picture). The model on the right side highlights all important amino acids for succinyl-CoA binding, in red (RcA)/blue (vAlaS) sticks. Some deviations of vAlaS are observed at position 143, 156, and 158. Asterisks indicate amino acids from the second monomer.

**Figure S4.**

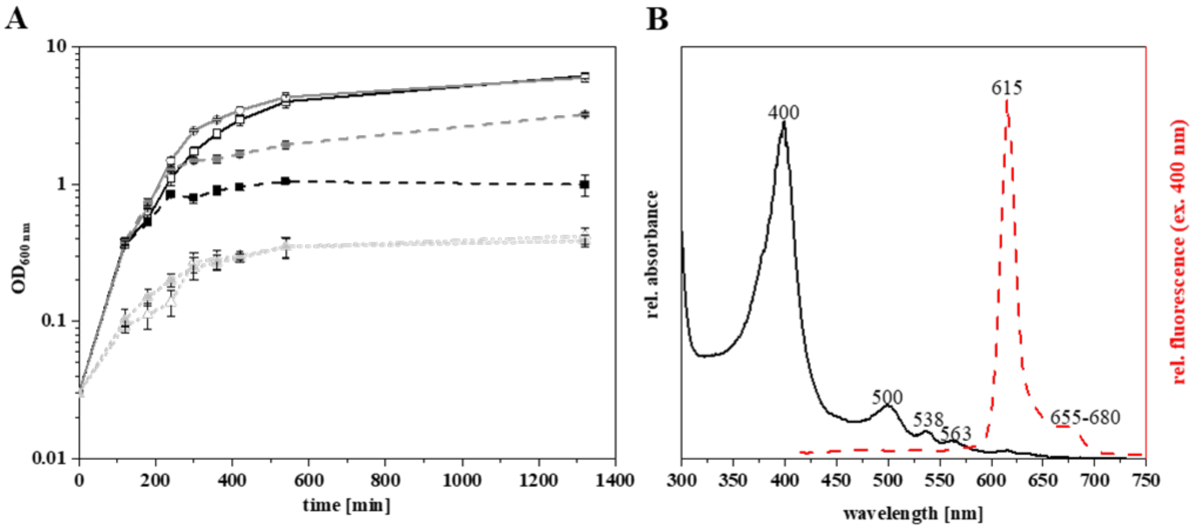


**Fig. S4. A.** Functional complementation of ALA-auxotrophic *E. coli* strain ST18 via plasmid-based complementation. *E. coli* growth was measured in the presence (filled symbol) or absence (empty symbol) of gene expression inductor IPTG for *tac* promotor controlled C4 or C5 pathway genes in comparison to the empty vector control. GtrR (black squares), RcA (grey circle), empty vector (light grey triangle). Mean values of three independent replicates are shown with standard deviation. **B.** Viral AlaS leads to porphyrin accumulation. Cell lysate absorption (black, solid line) and fluorescence (red, dashed line) measurement, show Soret and Q band, while fluorescence excitation at 400 nm show Stokes shift, indicating the presence of porphyrins in the cell lysate.

**Figure S5.**

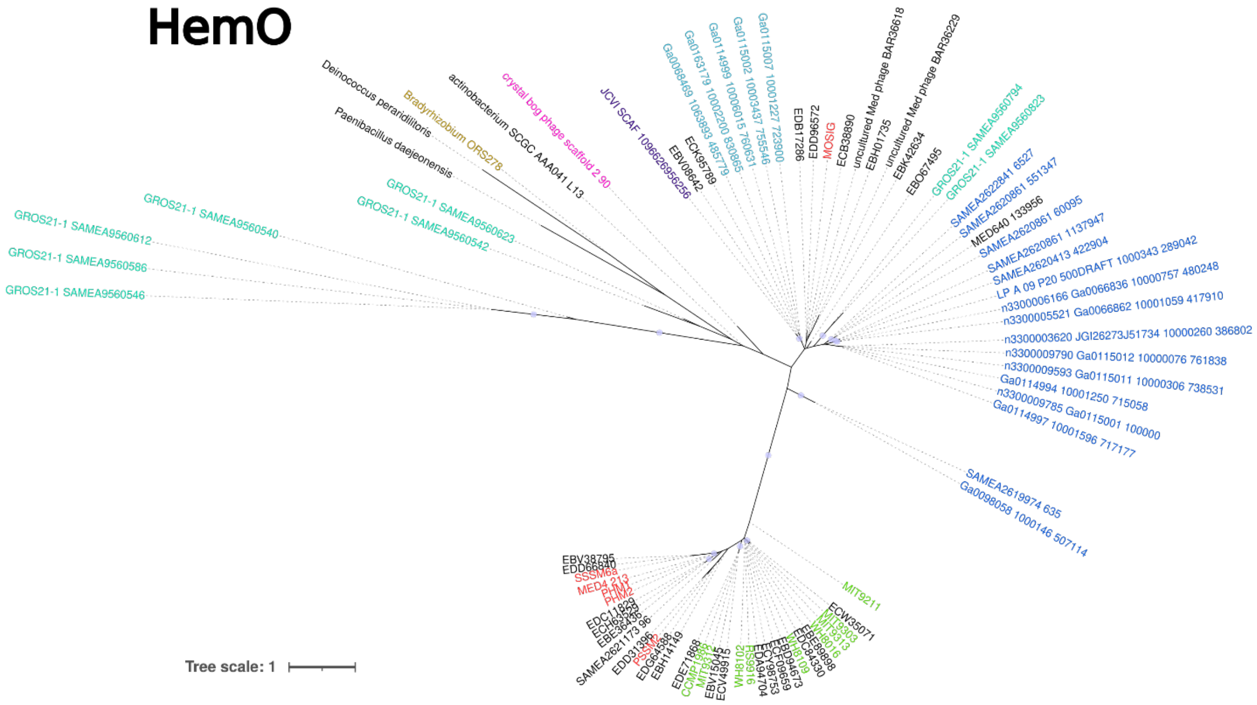


**Figure S5: Phylogenetic analyses of HemO protein sequences including sequences derived from contigs containing v*alaS*.** Maximum likelihood phylogenetic trees for HemO. Cyanobacterial strains are marked in light green; cultured phages are coloured red; black names denote noncyanobacteria and metagenomic contigs of uncertain origin. The *Bradyrhizobium* bacteria containing the three gene-cassette is marked in gold. The CB_2 phage chosen for experimental characterization is coloured pink. The previously characterized *hemO_pcyX* cassette (Ledermann et al., 2016) is marked in purple. Metagenomically retrieved contigs form this project are colour coded according to the bars in Figure 2. Circles represent bootstrap values >0.9. The scale bar indicates the average number of amino-acid substitutions per site.

**Figure S6.**

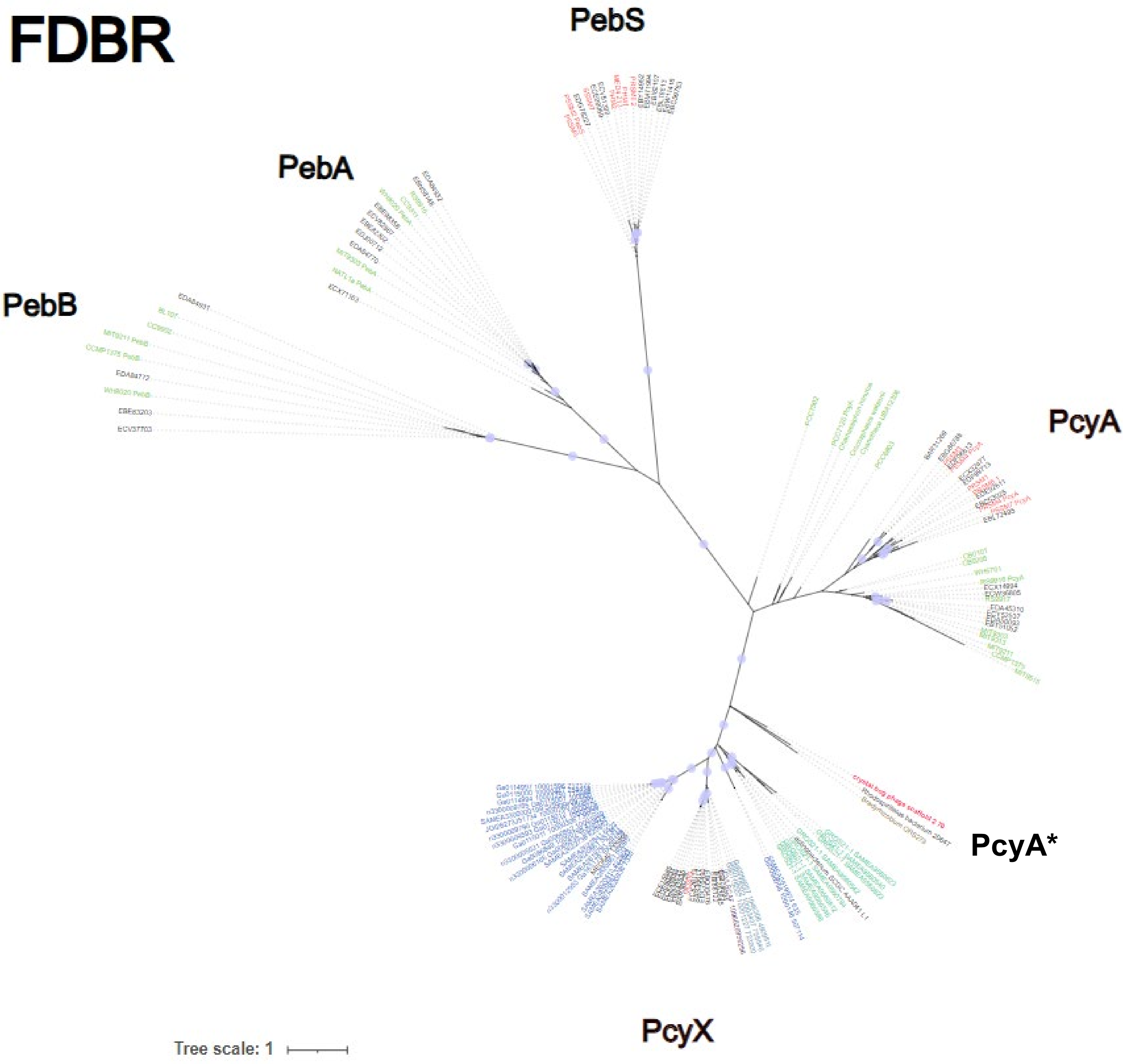


**Figure S6: Phylogenetic analyses of ferredoxin dependent bilin reductases (FDBRs) including sequences from contigs containing v*alaS*.** Maximum likelihood phylogenetic tree for FDBRs. Cyanobacterial strains are marked in light green; cultured phages are coloured red; black names denote noncyanbacteria and metagenomic contigs of uncertain origin. The *Bradyrhizobium* bacteria containing the three gene-cassette is marked in gold. The CB_2 phage chosen for experimental characterization is coloured pink. The previously characterized cassette (Ledermann et al., 2016) is marked in purple. Metagenomically retrieved contigs from this project are colour coded according to the bars in Figure 2. Circles represent bootstrap values >0.9. The scale bar indicates the average number of amino-acid substitutions per site. PcyA* marks a subgroup of PcyA enzymes including the one investigated in this study as well as the one from *Bradyrhizobium* sp. ORS278 investigated earlier (Jaubert et al., 2007; Ledermann et al., 2018).
